# ALS mutations disrupt self-association between the Ubiquilin Sti1 hydrophobic groove and internal placeholder sequences

**DOI:** 10.1101/2024.07.10.602902

**Authors:** Joan Onwunma, Saeed Binsabaan, Shawn P. Allen, Banumathi Sankaran, Matthew L. Wohlever

**Author notes:** These authors contributed equally (Joan Onwunma, Saeed Binsabaan, Shawn P. Allen).

## Abstract

Ubiquilins are molecular chaperones that play multifaceted roles in proteostasis, with point mutations in UBQLN2 leading to altered phase separation properties and Amyotrophic Lateral Sclerosis (ALS). Our mechanistic understanding of this essential process has been hindered by a lack of structural information on the Sti1 domain, which is essential for Ubiquilin chaperone activity and phase separation. Here, we present the first crystal structure of a Ubiquilin family Sti1 domain bound to a transmembrane domain (TMD) and show that ALS mutations disrupt the Sti1-TMD interaction. We then demonstrate that Ubiquilins contain multiple conserved, internal sequences that bind to the Sti1 domain, including the PXX region which is a hotspot for ALS mutations. We propose that these placeholder sequences prevent solvent exposure of the Sti1 hydrophobic groove and contribute to the multivalency that drives Ubiquilin phase separation. Together, this work provides a new paradigm for understanding how Sti1 domains modulate Ubiquilin chaperone activity and phase separation and offer insights into the molecular basis of ALS pathogenesis.

## Introduction

Amyotrophic lateral sclerosis (ALS) is a fatal neurodegenerative disease characterized by the progressive degeneration of motor neurons^1^. A hallmark of ALS is dysregulated protein homeostasis, resulting in altered dynamics, composition, and material properties of biomolecular condensates^2–6^. The Ubiquilin family of proteins are key players in the cellular proteostasis network, facilitating the degradation of misfolded proteins, chaperoning uninserted membrane proteins, and interacting with other cellular components to regulate phase separation^7–11^. Humans have three widely expressed Ubiquilin paralogs, UBQLN1, UBLQN2, and UBLQN4, which are linked to numerous neurodegenerative diseases, with point mutations in UBQLN2 causing dominant, X-linked Amyotrophic Lateral Sclerosis (ALS)^12–15^. Despite decades of research, the mechanistic basis for how these mutations disrupt Ubiquilin function and lead to disease is unclear.

A defining feature of Ubiquilins is a propensity for phase separation. While the physiological role of phase separation remains opaque, growing evidence links ALS mutations with altered phase separation properties^16^. For example, histology studies of ALS patients and mouse models of neurodegenerative diseases show increased formation of UBQLN2 puncta^17–19^. Furthermore, *in vitro* studies show that many Ubiquilins have altered material properties^20,21^. Understanding how ALS mutations alter Ubiquilin phase separation properties remains an important goal for the field.

Ubiquilins are characterized by an N-terminal Ubiquitin Like (UBL) domain, two Sti1 domains, and a C-terminal Ubiquitin Associated (UBA) domain, separated by long stretches of intrinsically disordered regions (IDRs) **(Figure 1A)**^7^. UBQLN2 also contains a unique PXX region, which is a hotspot for many ALS mutations. Recent studies have identified additional ALS mutations in both UBQLN2 Sti1 domains, which remain poorly characterized^17^. Interestingly, the Sti1-II domain is essential for UBQLN2 condensate formation, but how this domain contributes to phase separation is unclear^22^.

**Figure 1:**
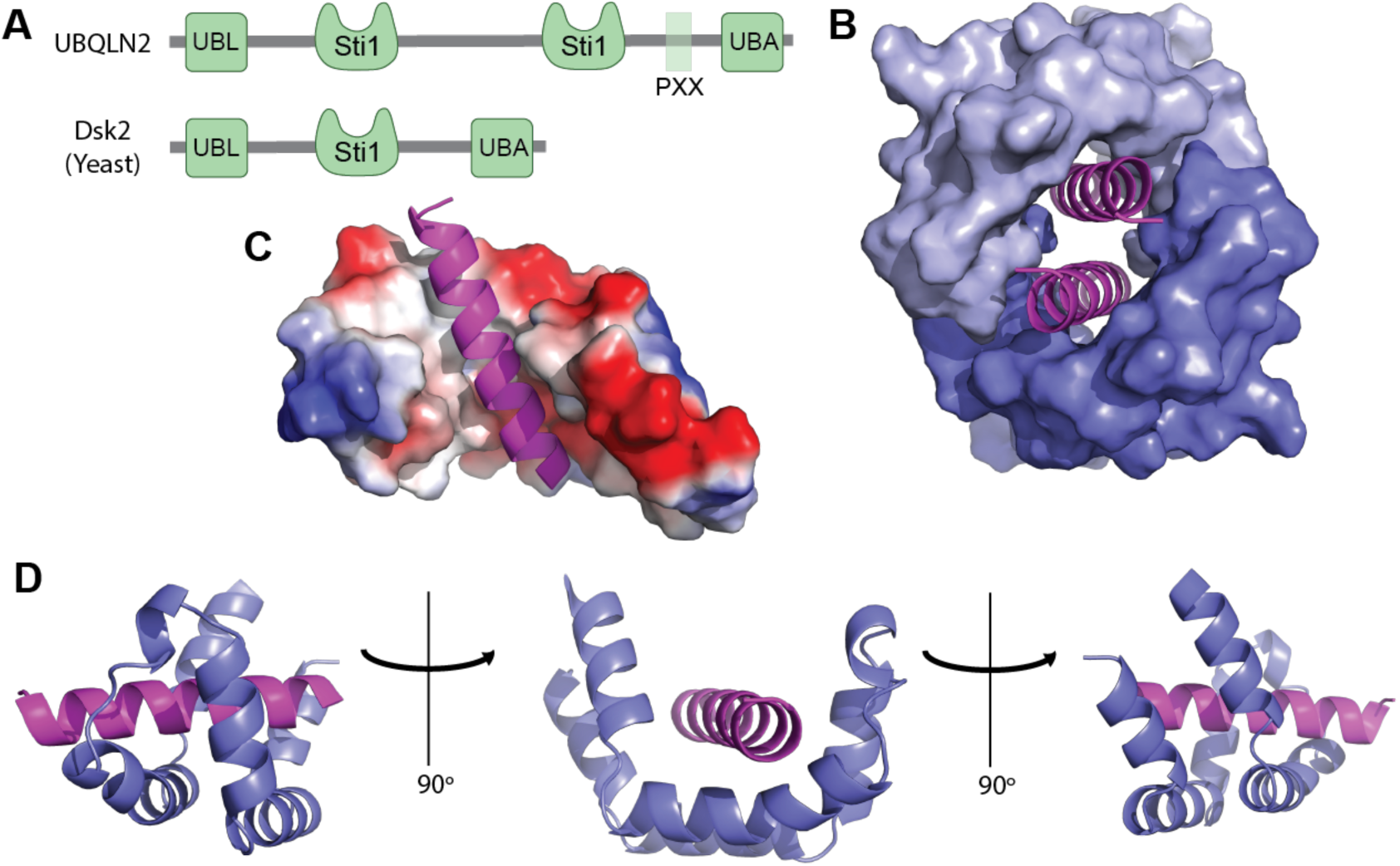
Crystal structure of the Dsk2 Sti1 domain bound to a TMD. A) Domain diagram of UBQLN2 and Dsk2. B) Dimerization of the Dsk2 Sti1 domains forms a hydrophobic cavity that fully encapsulates two TMDs. Individual subunits are shown as a surface and TMDs are shown as cartoons. FrzS domains omitted for clarity (PDB: 9CKX). C) Surface representation of Sti1 domain, colored by electrostatics. Note how the TMD binds in a hydrophobic groove. D) Cartoon representation of a single Sti1 domain. The TMD is colored magenta and the Sti1 domain is colored slate.

Despite an initial characterization as proteasomal shuttle factors that facilitate degradation of ubiquitinated client proteins^23–26^, Ubiquilins also have intrinsic chaperone activity, which is comparatively understudied^27,28^. Previous work has shown that Ubiquilin Sti1 domains can directly bind to moderately hydrophobic transmembrane domains (TMDs) a motif that is particularly enriched among mitochondrial membrane proteins^29,30^. While the Sti1 domains are rich in methionine residues, a hallmark of many TMD binding proteins^31–33^, the molecular basis for selectively binding moderately hydrophobic TMDs is unclear. Sti1 domains are a common motif in proteostasis and can be broadly categorized into co-chaperone and adaptors of the Ubiquitin Proteasome System^34,35^. There is little structural data on the proteasome adaptor family Sti1 domains, which includes Ubiquilins. While several modeling studies, including AlphaFold, have predicted the rough shape of the Ubiquilin Sti1 domain^34,36^, these predictions have low confidence. The lack of experimental structural data on Sti1 domain chaperone activity is thus a major roadblock in the field.

Here, we present the first crystal structure of the Ubiquilin family Sti1 domain bound to a TMD. Structure-function analysis shows that ALS causing mutations disrupt the Sti1 hydrophobic groove and TMD binding. We also developed a novel barcoded binding assay to discover that Ubiquilin Sti1 domains preferentially bind to hydrophobic sequences with low helicity, a property shared by multiple sequences within Ubiquilins. We then demonstrate that Ubiquilins contain multiple conserved, internal sequences that bind to the Sti1 hydrophobic groove, which we term placeholder sequences. Importantly, these placeholder sequences include the PXX motif in UBQLN2, which is a hotspot for ALS mutations that modulate Ubiquilin phase separation. We propose that the Sti1-placeholder interaction prevents solvent exposure of the Sti1 hydrophobic groove and contributes to the multivalency that drives Ubiquilin phase separation, thereby explaining why ALS causing mutations alter phase separation properties.

## Results

### Crystal structure of the Sti1 domain bound to a TMD

Ubiquilin Sti1 domains are flexible and have short-lived interactions with substrates, making them challenging targets for traditional structural biology approaches^29,34^. To stabilize the Sti1-TMD interaction we generated a fusion construct consisting of the Sti1 domain from *M. bicuspidata* Dsk2, a variant of the VAMP2 TMD, and the receiver domain from *Myxococcus xanthus* social motility protein FrzS as a crystallization scaffold^37^. FrzS was chosen because it robustly crystallizes at high resolution and has the N and C-termini positioned to allow for optimal interaction between the TMD and Sti1 domain^38^. The resulting construct formed crystals within three days at 4° C. We solved the crystal structure via molecular replacement with the FrzS domain with an overall resolution of 1.98 Å.

Our structure shows a domain swapped dimer, with the TMD of subunit 1 binding to the Sti1 domain of subunit 2 **(figure S1A)**. Two Sti1 domains stack head-to-head to create a fully enclosed hydrophobic cavity capable of chaperoning two TMDs **(Figure 1 & S1)**. The presence of multiple TMDs within the hydrophobic groove is reminiscent of the crystal structure of the TMD chaperone and targeting factor GET3^39^. The dimer is unlikely to be an artifact of crystallization as the two FrzS domains do not contact each other in the asymmetric unit and the crystallization construct runs as an oligomer on size exclusion chromatography **(Figure S1)**. The dimerization of the Sti1 domain is also thermodynamically appealing as it ensures that no part of the hydrophobic TMD is solvent exposed. Interestingly, the Sti1-II domain is essential for UBQLN2 dimerization^40^. Additionally, NMR studies on *S. cerevisiae* Dsk2 support Sti1-Sti1 interactions, suggesting that dimerization of the Sti1 domain may be a key feature of the Ubiquilin function^41–43^.

Each Sti1 monomer is composed of 6 α-helices that assemble into a helical-hand or U-shape. Each of the sides of the helical-hand contain two α-helices connected by short loops with ∼90° turns **(Figure 1D)**. The hydrophobic groove is enriched in methionine residues **(Figure S1D)**, a common feature of many membrane protein chaperones^31,44^. The overall shape is roughly similar to the AlphaFold2 prediction, however there are numerous differences in the details of TMD packing with the hydrophobic groove, highlighting the importance of experimental structural biology approaches **(Figure S1E)**. The TMD binds the bottom of the hydrophobic groove and buries ∼2970 Å^2^ of hydrophobic surface area with each Sti1 monomer. Key residues in the hydrophobic groove that contact the TMD include L190, M193, A207, M210, L211, M214, M218, M225, M228, and M229, which correspond to residues 30, 33, 47, 50, 51, 54, 58, 65, 68, and 69 in the crystallization construct. We conclude that the Sti1 domain forms a hydrophobic groove capable of forming head-to-head dimers that can fully encapsulate two TMDs.

### ALS Mutations disrupt the Sti1 hydrophobic groove

To gain a better understanding of how ALS mutations in the Sti1 domain affect Ubiquilin function, we performed a sequence alignment of the *M. bicuspidata* Sti1 domain against the Sti1-I or Sti1-II domains from UBLQN1, 2, and 4 **(Figure S2)**. The resulting sequence alignment was then mapped onto our crystal structure and the position of UBQLN2 ALS mutations was analyzed. Our structure predicts that the Q425R and M446R mutations will disrupt the hydrophobic groove via insertion of a charged residue **(Figure 2A & B)**. Notably, the AlphaFold prediction of UBQLN2 yields similar results.

**Figure 2:**
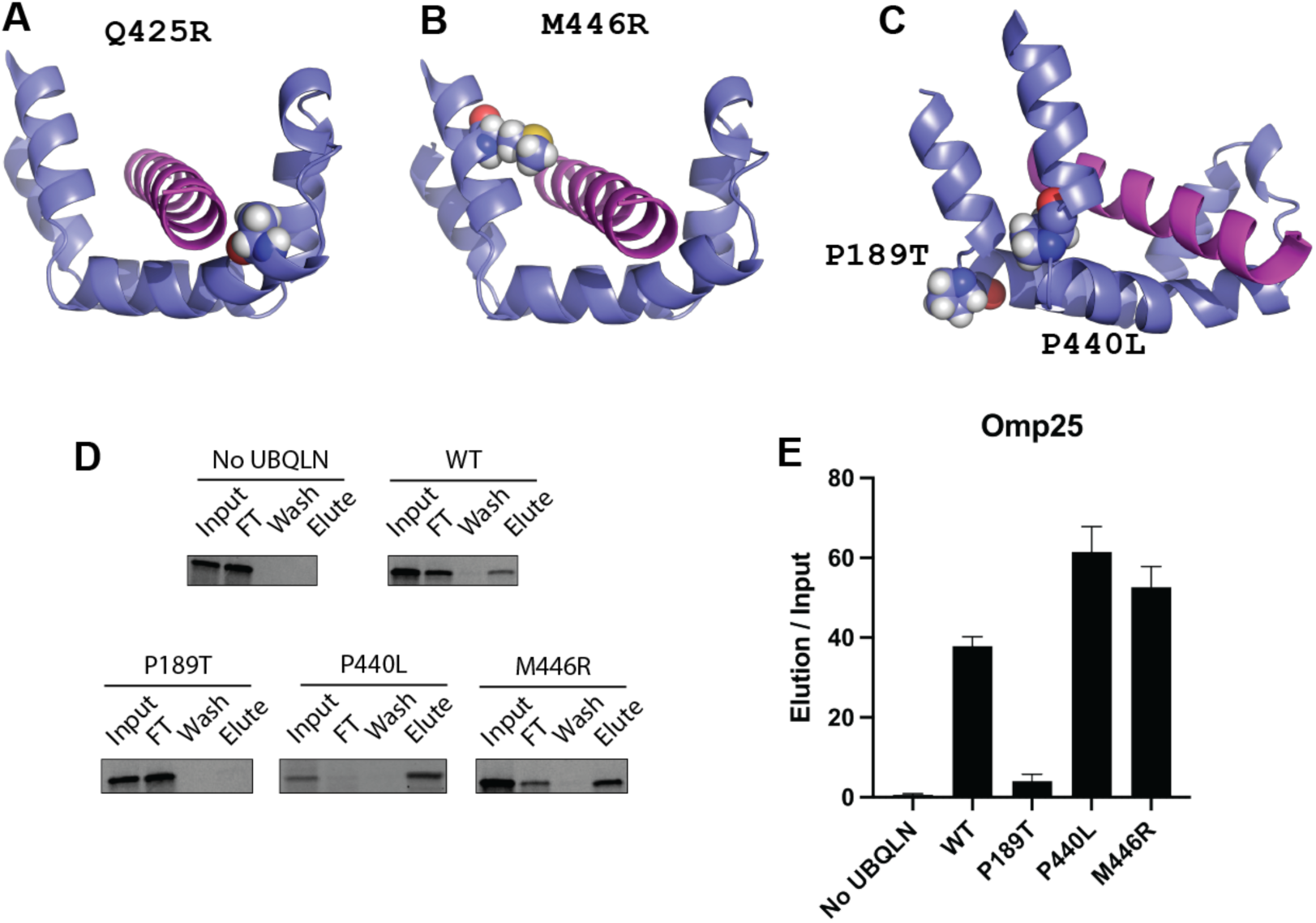
UBQLN2 ALS mutations disrupt the Sti1 hydrophobic groove. A) The UBQLN2 sequence was mapped onto the Dsk2 structure. The UBQLN2 Q425R mutation, which causes ALS, is predicted to insert a charged residue into the Sti1 hydrophobic groove. B) The UBLQN2 M446R mutation is also predicted to insert a charged residue into the Sti1 hydrophobic groove. C) Residues P189 and P440 (spheres) in UBQLN2 are essential for forming the ∼90° turns between α-helices in the helical hand motif. Mutation of these highly conserved residues is predicted to disrupt formation of the hydrophobic groove. D) Omp25 binds to Sti1-1. The ^35^S labeled Omp25 TMD was produced by PURExpress IVT in the presence of recombinant Flag_3x_-UBQLN2 variants and binding was assayed by anti-Flag immunoprecipitation. E) Quantification of data in D, error bars are standard error of the mean, N ≥ 3.

The P189T and P440L mutations also occur in the UBQLN2 Sti1 domain but are not predicted to contact the TMD **(Figure 2C)**. Our analysis shows that the P189T and P440L mutations occur in short loops with a ∼90° turn between sides of the helical-hand. The prolines and preceding residues have Phi-Psi angles that are uniquely favored by proline^45^. We propose that mutation of these highly conserved proline residues will disrupt folding of the Sti1 domain.

To test these predictions, we used the PURExpress In Vitro Transcription/Translation (IVT) system to produce ^35^S labeled substrates in a detergent free environment^46^. The PURExpress reaction was supplemented with physiological concentrations of recombinantly purified flag-tagged UBQLN2 variants. As the PURExpress system contains no other chaperones, membrane, or detergent, binding of the membrane protein to Ubiquilins can be assessed by a simple immunoprecipitation (IP). We chose the mitochondrial membrane protein Omp25 as our model substrate as it had been previously shown to bind to UBQLN1 Sti1 domains^29^ .

We observed robust binding of Omp25 to WT UBQLN2 and no binding in the absence of Ubiquilins. Our structure predicts that the P189T and P440L mutations will be the most disruptive to substrate binding. Interestingly, we observed that the P189T mutation leads to a 90% decrease in substrate binding whereas the P440L mutation has no effect. As the P189T mutation is in the Sti1-1 domain and the P440L mutation is in the Sti1-2 domain, this suggests that Omp25 binds exclusively to Sti1-1. Consistent with this interpretation, the M446R mutation in the Sti1-2 domain also has no effect on Omp25 binding. While our structure clearly predicts that the Q425R, P440L, and M446R mutations should disrupt the hydrophobic groove of Sti1-2, testing this model will require a substrate that binds to Sti1-2. We conclude that Omp25 binds to the Sti1-1 domain and that the ALS causing P189T mutation disrupts Omp25 binding.

### Development of a barcoded binding assay to study substrate binding to the Sti1 domain

It was previously proposed that Ubiquilins selectively bind to moderately hydrophobic TMDs^29^. Our crystal structure shows a highly hydrophobic groove and cannot easily explain a preference for binding moderately hydrophobic TMDs. We therefore decided to rigorously test this model by developing a competitive binding assay.

Substrate was again produced via the PURExpress IVT system supplemented with physiological concentrations of recombinantly purified flag-tagged Ubiquilin. To better mimic the complexity of the cellular environment, we sought to simultaneously present Ubiquilins with multiple closely related substrates. We developed a pool of six model substrates based on the TMD of Omp25, a mitochondrial tail anchored (TA) protein and known Ubiquilin substrate^29,47^. The hydrophobicity of the substrate TMDs, as assessed by Grand Average of Hydrophobicity (GRAVY) scores^48^, was systematically varied by mutating residues to Alanine or Leucine. To differentiate substrates by SDS PAGE, a variable number of SH3 domains were added to serve as a “barcode” **(Figure 3A)**. The SH3 domain was chosen because it is small, well-folded, and contains no methionine residues which would bias the ^35^S signal intensity^49^. After testing the expression of each plasmid individually, the six plasmids for a substrate series were mixed together such that expression of each model substrate was roughly equal. Ubiquilin-substrate complexes were purified by anti-Flag IP and binding was calculated by comparing the amount of substrate in the elution and input fractions.

**Figure 3:**
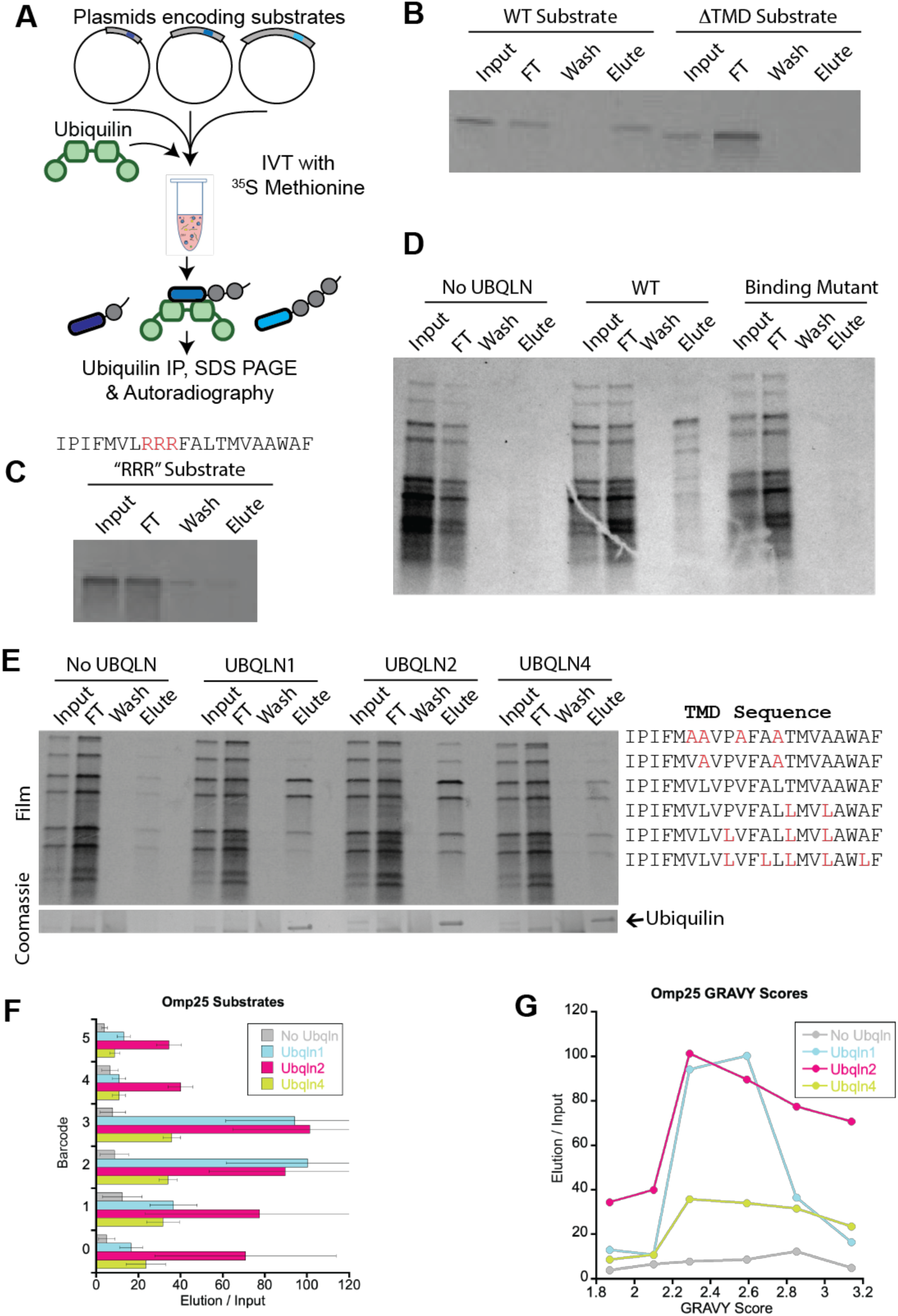
Development of barcoded binding assay to study Ubiquilin-substrate binding. A) Overview of the barcoded binding assay. B) There is no biding to the barcode 3 construct when the TMD is deleted. C) Mutation of the Omp25 TMD with three consecutive Arginine residues is sufficient to disrupt binding to Ubiquilin. D) Substrate binding depends on interaction with the Sti1 domains. Binding assay with the Omp25 substrate series with no Ubiquilin control, WT UBQLN1, and the UBQLN1 binding mutant. The binding mutant contains two mutations in each Sti1 domain that are predicted to disrupt the hydrophobic groove (M186D, M228D, M394D, M454D). E) Barcoded binding assay with the Omp25 substrate series. The wild type Omp25 TMD has a barcode of 3. Residues were mutated to either Leucine or Alanine to change TMD hydrophobicity. F) Quantification of E, organized by barcode. Error bars are SEM. A value of 100 corresponds to equal intensity of input and elution bands. G) Quantification of E, organized by TMD hydrophobicity (GRAVY). Error bars are omitted for clarity.

Control experiments show that binding is dependent on the Sti1-TMD interaction. Deletion of the TMD shows interaction that is equivalent to our no Ubiquilin control, arguing that there is no interaction between Ubiquilins and the SH3 domains **(Figure 3B)**. Similarly, the previously characterized RRR TMD mutant^29^, which has three Arginine mutations in the middle of the TMD, disrupts substrate binding **(Figure 3C)**. Finally, we used our crystal structure and AlphaFold model to generate the UBLQN1 binding mutant, which contains two mutations in each Sti1 domain that are predicted to disrupt substrate binding. We observed no binding of any of the Omp25 substrates to the binding mutant, confirming that binding is dependent on the Sti1-TMD interaction **(Figure 3D)**.

### Ubiquilin paralogs show substantial overlap in substrate binding specificity

To test the substrate binding preference of Ubiquilins, we constructed a total of four substrate series, two based on the mitochondrial tail-anchored proteins Omp25 **(Figure 3E-G)** and Tom5 **(Figure S4)** and two based on the ER tail-anchored proteins Sec61β **(Figure S3)** and Vamp2 **(Figures S5)**. We chose tail-anchored proteins as our model substrates because they are verified Ubiquilin substrates and the single TMD simplifies data analysis. The wild type TMD sequence was placed in the middle of the series with a barcode of either 2 or 3 and the hydrophobicity of the TMD within a substrate series was varied by mutating residues to Leucine or Alanine such that the GRAVY score shifted by 0.2-0.3 for each substrate. The wild type sequences of the four series have a broad range of hydrophobicity values, with GRAVY scores ranging from 1.61 to 3.29.

The Omp25 and Sec61β substrate series showed that Ubiquilins bound preferentially to TMDs with moderate hydrophobicity **(Figure 3 & S3)**. This trend was conserved between UBQLN1, 2, & 4. However, the TMD hydrophobicity that elicited maximum Ubiquilin binding differed between the two substrate series. The Tom5 substrate series was the least hydrophobic substrate series **(Figure S4)**. It had the lowest overall level of binding. It still showed a preference for moderately hydrophobic TMDs, but the trend was much less pronounced than with Omp25 and Sec61β. The Vamp2 substrate series was the most hydrophobic series and, unexpectedly, showed the highest overall level of substrate binding with no clear up and down pattern **(Figure S5)**.

Overall, our results show that UBQLN1, 2, & 4 show a general trend for binding to TMDs with moderate hydrophobicity. However, the TMD GRAVY score that elicited maximum substrate binding varied based on the substrate series. This suggests that TMD GRAVY score alone fails to fully capture the biophysical parameters that govern substrate binding.

We observed modest, but reproducible differences in the substrate binding preferences of the Ubiquilin paralogs. UBQLN4 consistently had the lowest levels of substrate binding, with the results being most pronounced in the Omp25 **(Figure 3)** and Tom5 **(Figure S4)** substrate series. Within the Vamp2 substrate series, which was the most hydrophobic series, there were no statistically significant differences in substrate binding **(Figure S5)**.

UBQLN1 and UBQLN2 showed roughly equal levels of substrate binding, with the exception of the least hydrophobic substrate series, Tom5. However, as TMD hydrophobicity reached the more extreme values, UBQLN1 binding dropped close to background levels whereas UBQLN2 binding remained significantly above background levels. This trend is clearest in the Omp25 substrate series **(Figure 3)**, but is also observed with Sec61β **(Figure S3)** and Tom5 **(Figure S4)**. Overall, we conclude that there is substantial redundancy in the substrate binding preferences of UBQLN1, 2, & 4, but UBQLN2 has the most robust substrate binding and the broadest substrate range whereas UBQLN4 shows the lowest levels of substrate binding.

### Ubiquilin Sti1 domains selectively bind hydrophobic substrates with low helical propensity

Although Ubiquilin binding within each substrate series showed maximal binding to TMDs with moderate hydrophobicity, the GRAVY score that elicited maximal binding differed between each substrate series. When all four substrate series are analyzed together, there is no discernable preference for moderately hydrophobic TMDs **(Figure 4A)**. This indicates that substrate GRAVY score alone is a poor predictor of Ubiquilin binding.

**Figure 4:**
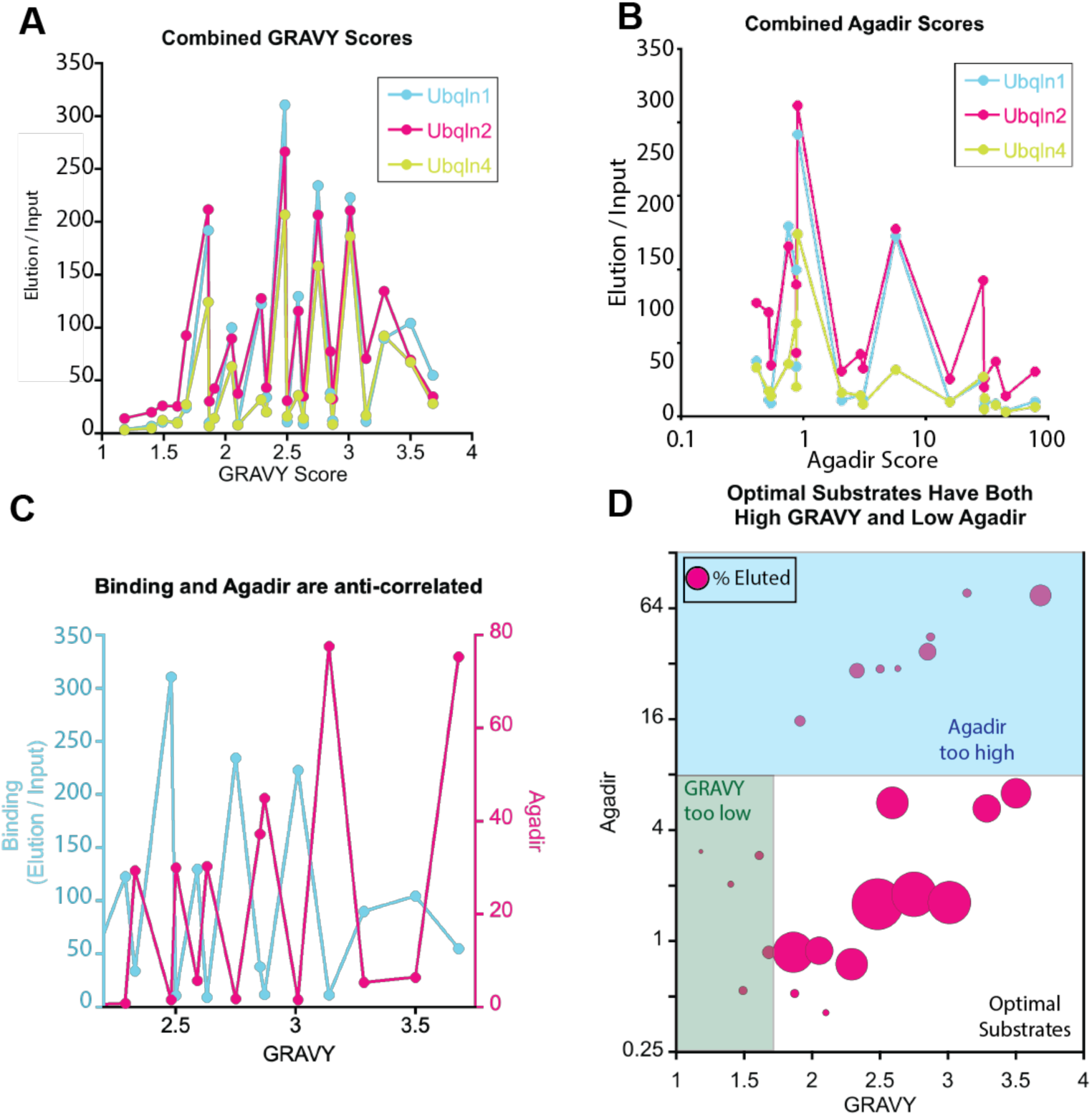
Barcoded binding assay shows that Ubiquilins bind to hydrophobic, disordered regions. A) Combined data from all four substrate series does not show a clear pattern between TMD GRAVY score and Ubiquilin binding. B) Combined data from all four substrate series does not show a clear pattern between TMD Agadir score and Ubiquilin binding. C) Combined data from all four substrate series shows that there is a strong anti-correlation between substrate binding and Agadir scores. D) Bubble chart shows that optimal Ubiquilin substrates have both a high GRAVY score and a low Agadir score.

Previous studies had demonstrated that ER TA proteins have higher TMD hydrophobicity and higher helical propensity than mitochondrial TA proteins^30^. As Ubiquilins were shown to preferentially bind to mitochondrial membrane proteins^10,29^, we asked if helical propensity also governed Ubiquilin substrate binding. The helical propensity of each TMD was calculated using the Agadir score^50–52^, with a high Agadir score corresponding to a high helical propensity. There was no obvious correlation between Agadir score and Ubiquilin binding **(Figure 4B)**.

We next asked if a combination of TMD GRAVY and Agadir scores are a better predictor of Ubiquilin binding. When plotted together, we observed a striking anti-correlation between substrate binding and Agadir score **(Figure 4C)**. When plotted as a bubble chart, we observed that optimal Ubiquilin substrates are hydrophobic TMDs with a low helical propensity **(Figure 4D)**. As many hydrophobic residues also have high helical propensity, this could account for the previous observation that Ubiquilins preferentially bind to moderately hydrophobic TMDs. We conclude that Ubiquilins have no inherent defect in binding to highly hydrophobic TMDs and substrate binding requires a combination of sufficient hydrophobicity and low helical propensity.

### Internal Ubiquilin Sequences bind to the Sti1 hydrophobic groove

Having determined the biophysical parameters that govern substrate binding to the Sti1 domains, we next asked what prevents the thermodynamically unfavorable solvent exposure of the Sti1 hydrophobic groove in the absence of substrate. Acharya et al. recently identified several sequences within *S. cerevisiae* Dsk2 that interact with the Sti1 domain^53^. Interestingly, these placeholder sequences are in the intrinsically disordered regions of Dsk2. Given our results that Ubiquilin Sti1 domains preferentially bind to regions of low helicity and the observation that Ubiquilins have several long stretches of disordered sequence, we hypothesized that Ubiquilins contain similar internal sequences capable of self-interaction with the Sti1 domain.

To test this hypothesis, we first asked if the placeholder sequences identified in *S. cerevisiae* Dsk2 are conserved in *H. sapiens* UBQLN2. Despite the significant size difference between UBQLN2 (624 residues) and Dsk2 (373 residues), we observed moderate conservation of these sequences **(Figure S6)**. We will refer to these putative placeholder sequences as PH1, 2, and 3. An analysis of the UBQLN2 AlphaFold-predicted structures shows that putative placeholders have a modest helical content and border unstructured regions **(Figure 5A & S7)**.

**Figure 5:**
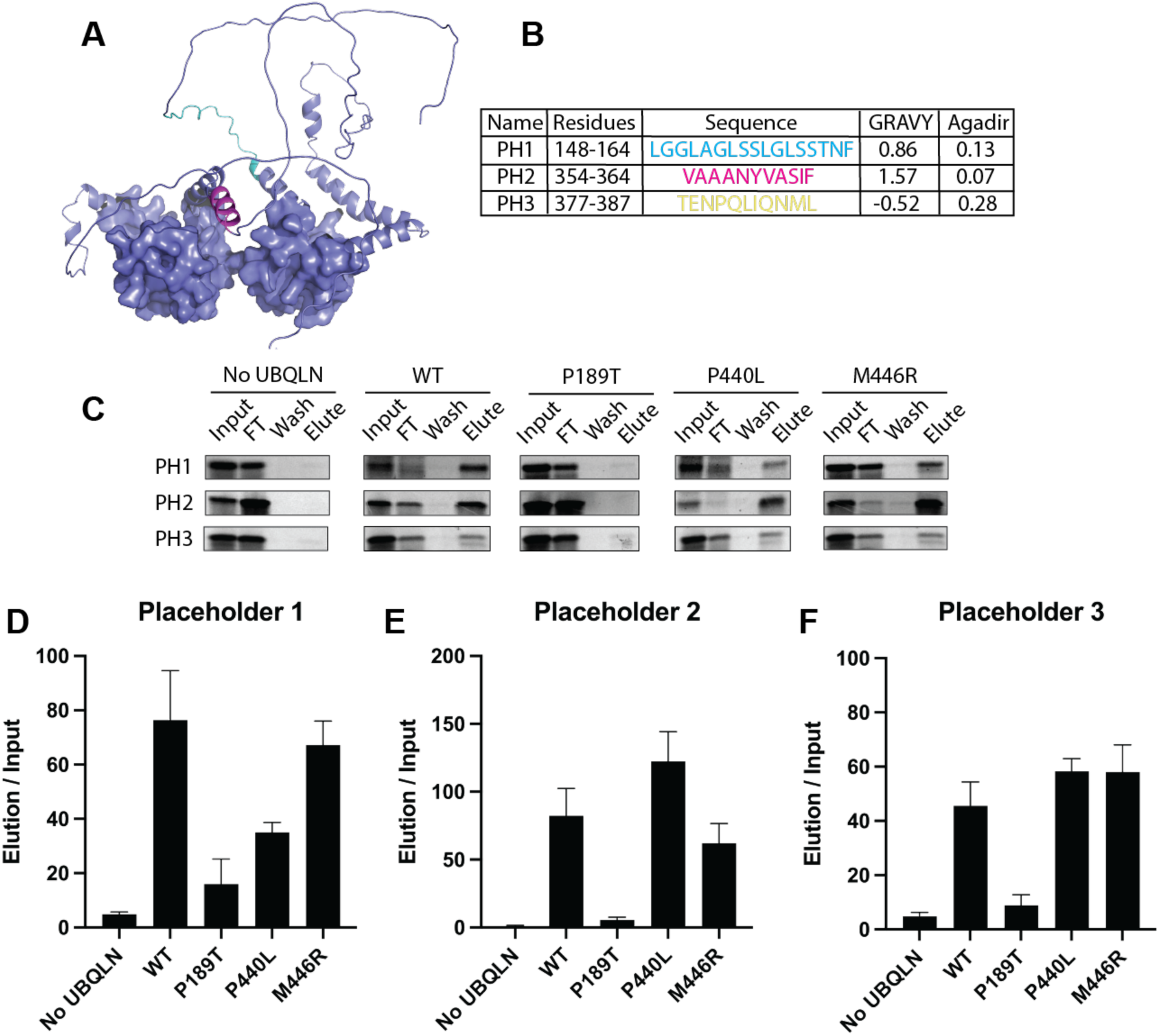
Internal placeholder sequences bind to the Sti1 hydrophobic groove. A) AlphaFold model of UBQLN2 shows that placeholder sequences 1 and 2 have modest helical content and are adjacent to unstructured regions. PH1 and PH2 are colored cyan and magenta respectively. Sti1 domains are shown as surfaces. UBL and UBA domains are omitted for clarity. B) Summary of biophysical properties for UBLQN2 placeholder sequences. C) Representative results of placeholder 1, 2, or 3 binding to UBQLN2 variants. D-F) Quantification of C, error bars are standard error of the mean, N ≥ 3.

We next examined the biophysical properties of PH1, 2, & 3 and observed that the putative placeholder sequences all have low Agadir scores. Our model predicts that placeholder sequences must have suboptimal binding characteristics, otherwise the high effective concentration of the placeholder sequence will prevent exogenous sequences from binding to the Sti1 domain. Consistent with our model PH1 and PH2 are generally hydrophobic, but the GRAVY scores fall outside the optimal range for a true substrate **(Figure 5B)**. By contrast, the overall sequence of PH3 is not hydrophobic and has a negative GRAVY score. Closer examination of the AlphaFold2 model reveals that PH3 forms a short, broken amphipathic helix with the hydrophobic face perfectly positioned to interact with the hydrophobic groove of Sti1 **(Figure S7)**.

To test if PH1, 2, & 3 can bind to the Sti1 domain, we again used the PURExpress IVT system to synthesize the placeholder sequences in the presence of physiological concentrations of UBQLN2. We observed robust binding of all 3 placeholder sequences to WT UBQLN2 and minimal binding in the no Ubiquilin control, indicating that Sti1 domain interaction with placeholder sequences is conserved from yeast to humans **(Figure 5C)**.

To test how ALS mutations in the Sti1 domains affect placeholder binding, we repeated the assay with the P189T, P440L, and M446R UBQLN2 mutants **(Figure 5D)**. Unlike the results with the hydrophobic TMD Omp25, we observed that the P189T and P440L mutations each led to a partial loss of binding to PH1, suggesting that PH1 binds to both the Sti1-1 and Sti1-2 domains. The M446R mutation, which is predicted to be less disruptive than the P440L mutation, appeared to have a modest defect in binding that did not reach significance. Conversely, the PH2 and PH3 sequences bound exclusively to the Sti1-1 domain. We conclude that the interaction of Sti1 domains with internal placeholder sequences is conserved from yeast to humans, that both Sti1 domains in UBQLN2 are capable of interacting with internal placeholder sequences, and that ALS causing mutations within the Sti1 domains disrupt interaction with placeholder sequences.

### The PXX region predominantly binds to the Sti1-1 domain

Given our results that ALS mutations in the Sti1 domain disrupt the Sti1-placeholder interaction, we hypothesized that the UBQLN2 PXX region, an ALS mutation hotspot, also interacts with one of the Sti1 domains. Examination of the Agadir and GRAVY scores shows biophysical parameters consistent with the other placeholder sequences identified above **(Figure 6A)**. To test this hypothesis, we again used our PURExpress binding assay. Consistent with our hypothesis, we observed robust binding of the PXX region to WT UBQLN2 and minimal binding to the no Ubiquilin control **(Figure 6B)**. We then repeated the assay with the P189T, P440L, and M446R UBQLN2 mutants **(Figure 6C)**. We observed a strong decrease in binding with the P189T mutation and a modest decrease with the P440L and M446R mutations that did not reach significance. We conclude that the PXX region binds predominantly to the Sti1-1 domain.

**Figure 6:**
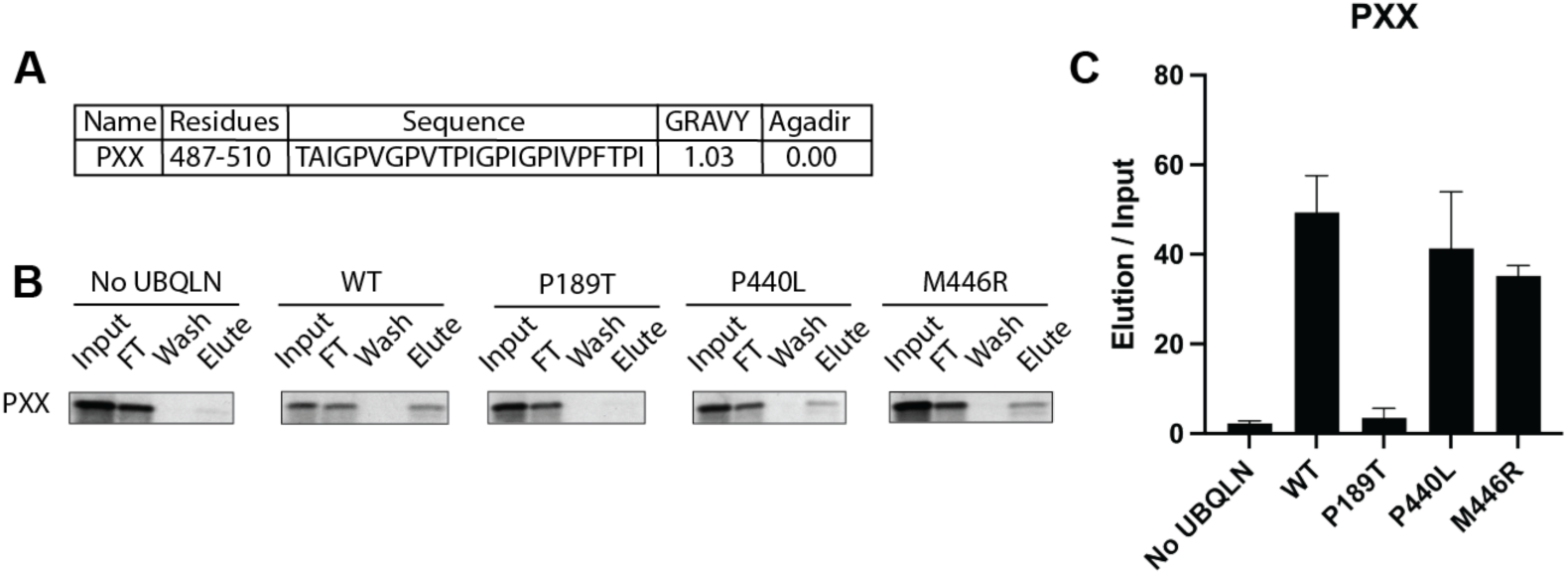
The PXX region predominantly binds to the Sti1-1 domain. A) The PXX region is hydrophobic (high GRAVY score) and has low helicity (low Agadir score), consistent with other placeholder sequences. B) Representative results of the PXX region binding to UBQLN2 variants. C) Quantification of B, error bars are standard error of the mean, N ≥ 3.

## Discussion

The Sti1 domains have been linked to numerous roles in Ubiquilin function and disease, such as chaperone activity, dimerization, phase separation, and ALS^7,17,29^. However, a clear mechanistic and structural explanation for these multifaceted roles remains elusive. Here, we address these key questions by solving the first crystal structure of a Ubiquilin Sti1 domain bound to a TMD and developing a novel barcoded binding assay to directly measure substrate interaction with the Sti1 domains. Our structure shows that the Sti1 domain forms a methionine rich hydrophobic groove. The barcoded binding assay demonstrates that Sti1 domains preferentially bind to hydrophobic sequences with low helicity, two motifs present throughout the long, disordered regions between annotated domains in Ubiquilins. Consistent with these results, we show that multiple conserved internal placeholder sequences, including the ALS associated PXX region, bind to the Sti1 domains. Finally, we demonstrate that multiple ALS-causing mutations interfere with the Sti1 hydrophobic groove, thereby disrupting binding to both TMDs and placeholder sequences. Together, these results lead to a new paradigm for understanding the role of the Sti1 domains in Ubiquilin phase separation and how this activity is dysregulated in ALS.

Our crystal structure shows a methionine rich hydrophobic groove, which readily explains Ubiquilin chaperone activity. In the absence of a bound TMD, the hydrophobic groove within each Sti1 domain will have thermodynamically unfavorable interactions with solvent. Many TMD chaperones therefore use internal placeholder sequences to occupy the binding sites and act as a substrate selectivity filter that is displaced upon encounter with a bona fide substrate^39,54,55^. Acharya et al. recently identified several sequences within *S. cerevisiae* Ubiquilin homolog Dsk2 that bind to the Sti1 domain^53^. Here, we demonstrate that similar placeholder sequences also exist in UBQLN2 and these sequences bind to both Sti1-I and Sti1-II domains **(Figure 7A & B)**. Our results suggest that the placeholder-Sti1 interaction is conserved from yeast to humans and is perhaps a general feature of Sti1 domain containing proteins.

**Figure 7:**
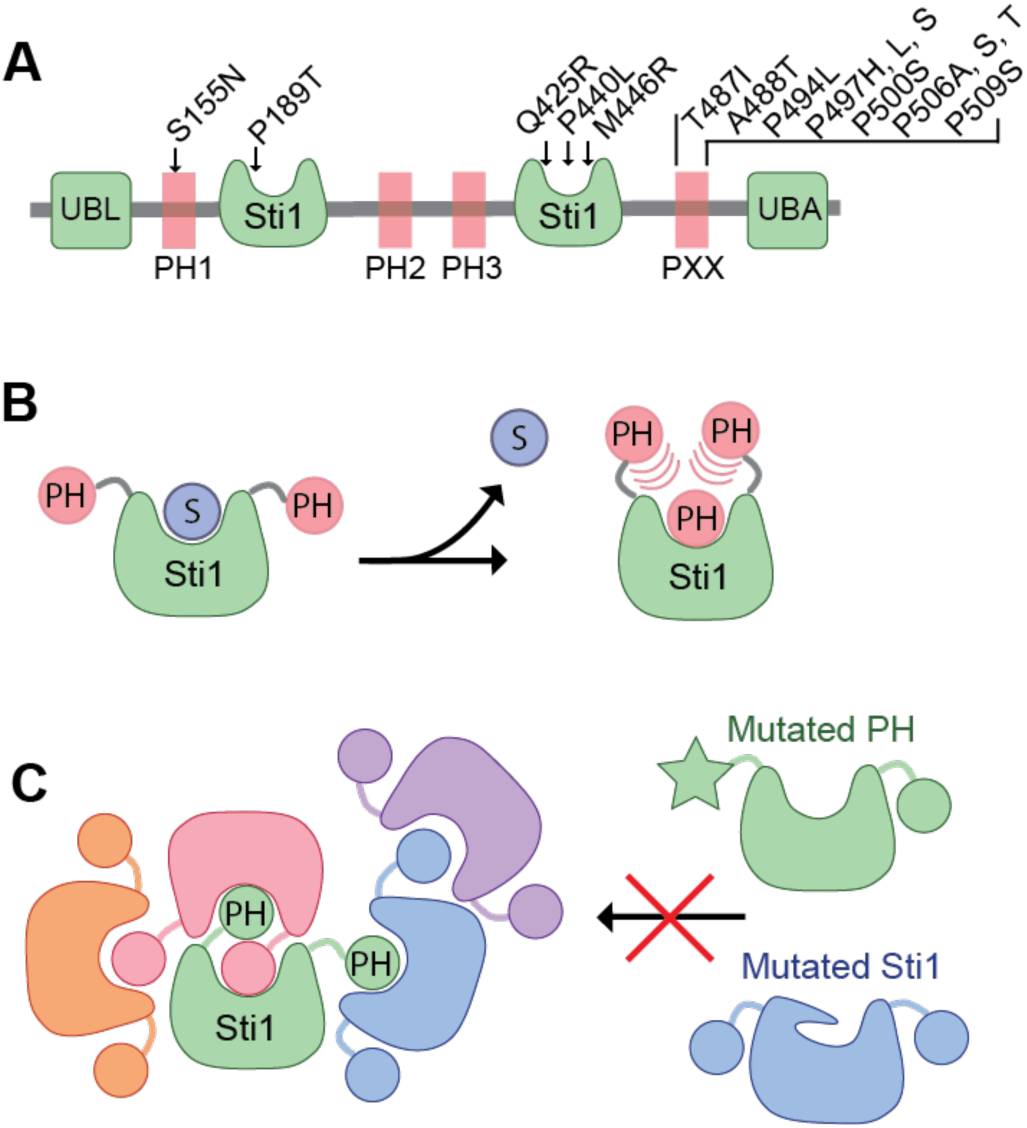
Proposed model for how ALS mutations lead to altered phase separation. A) Domain diagram of UBQLN2. The four identified placeholder sequences capable of interacting with the Sti1 domain are shown in red. ALS causing mutations within the placeholder and Sti1 domains are indicated. B) Hydrophobic substrates (S) bind to the Sti1 domain. In the absence of substrate, placeholders dynamically bind to the Sti1 domain to limit solvent exposure of the hydrophobic groove. C) Model for how intermolecular interactions between Sti1 domains and placeholders contribute to the multivalency required for phase separation. ALS mutations within the placeholder sequences or Sti1 domains are predicted to disrupt this interaction, leading to altered phase separation. For visual clarity each molecule of Ubiquilin is a different color and is represented with only one Sti1 domain and two placeholders.

The placeholder-Sti1 interaction has profound implications on Ubiquilin activity, particularly with substrate binding, Ubiquilin phase separation, and ALS. Based on the results of the barcoded binding assay, the biophysical properties of the placeholder sequences appear to be suboptimal for robust Sti1 domain interaction. However, the presence of multiple internal placeholder sequences will lead to a high effective concentration, such that the Sti1 domain is predominantly occupied by a placeholder sequence. For an exogenous substrate, such as a TMD, to stably bind to the Sti1 domain, it must have biophysical properties that allow it to outcompete the high effective concentration of the placeholder sequences. In this way, the placeholder sequences may serve as a selectivity filter to regulate substrate binding to the Sti1 domain.

A key driver of phase separation is multivalency. It is already well documented that the Ubiquilin UBL and UBA domains are important for self-interaction, however, it is less clear why the Sti1-II domain is essential for phase separation. We propose that the Sti1 domain contributes to phase separation in multiple ways. First, by the interaction of the Sti1 domain with multiple internal placeholder sequences and second, by Sti1 dimerization **(Figure 7C)**. The Sti1-placeholder interactions can occur intra- or intermolecularly, with intermolecular interactions becoming more prevalent as the total Ubiquilin concentration increases. The displacement of internal placeholder sequences by a bona fide substrate implies that substrate binding will modulate Ubiquilin phase separation by disrupting the Sti1-placeholder interaction. Directly testing this model will be important for solidifying our understanding Ubiquilin phase separation.

There are likely more placeholder sequences in Ubiquilins than those identified here. Indeed, paramagnetic spin labeling studies have shown that the UBQLN2 UBAA motif interacts with the Sti1-II domain^16^. Differences in the number and/or biophysical properties of the placeholder sequences could also explain the different phase separation properties of UBQLN1, UBQLN2, and UBLQN4 despite high sequence similarity in the Sti1 domains. Identification of all placeholder sequences will be important for fully modeling Ubiquilin phase separation

The dimerization of the Sti1 domains also contributes to multivalency. Similar to the Sti1-placeholder interactions, the dimerization of isolated Sti1 domains is likely transient. The extensive contacts between the TMDs in our crystal structure likely contributed to the stability of the dimer. Despite these caveats, the dimerization of the Sti1 domains is critical for Ubiquilin multivalency. Previous work has shown that the Sti1-II domain is required for UBQLN2 dimerization and recent NMR studies show that the Sti1 domain of yeast Dsk2 is involved in self-interaction^53^. The multiple contributions of the Sti1 domain to intermolecular interactions explains why the Sti1-II domain, but not the placeholder PXX region, is critical for Ubiquilin phase separation^22^ . Consistent with these results, Acharya et al. demonstrated that deletion of the Sti1 domain from the yeast Ubiquilin homolog Dsk2 has a larger effect on phase separation than co-deletion of three placeholder sequences^53^.

Many UBQLN2 ALS mutations lead to altered phase separation properties, but the mechanistic details of how this occurs have remained obscure. Our model now makes testable, structure-based predictions for how ALS mutations modulate Ubiquilin phase separation. Mutations in the Sti1 domain are predicted to disrupt the hydrophobic groove. Consistent with these predictions, we demonstrate that the pathogenic P189T and P440L mutations disrupt placeholder binding to the Sti1 domains. The interaction of the PXX region, with the Sti1 domains is particularly exciting as numerous ALS mutations are contained within this sequence **(Figure 7A)**.

Modulation of the Sti1-placeholder interaction therefore presents a new paradigm for therapeutic intervention in ALS patients with UBQLN2 mutations.

In summary, our findings provide a structural and mechanistic understanding of the Sti1 domains in Ubiquilin proteins, elucidating their roles in substrate binding, phase separation, and ALS pathogenesis. By revealing the conserved interplay between Sti1 domains and placeholder sequences, as well as the effects of ALS-related mutations, this study highlights the critical role of hydrophobic interactions in regulating Ubiquilin function. These insights lay the groundwork for future investigations into Ubiquilin-mediated cellular processes and potential therapeutic interventions for ALS and related disorders

## Materials and Methods

### Plasmid Construction

The bar-coded substrates have the general structure of 3xHA-Sec61β soluble domain-n x SH3-TMD-3F4, where n varies between 0 and 5 to generate the barcode. The Omp25 substrates were generated by gene synthesis (Azenta) and cloned into the PURExpress DHFR Control Vector (NEB) between the NdeI and NotI sites. Note that each SH3 domain had a unique DNA sequence and unique restriction enzyme sites between the SH3 domains and the TMD. Subsequent substrate series were generated by replacing the TMD sequence with a new TMD sequence generated by gene synthesis. New TMD sequences were restriction cloned between KpnI and BamHI. Ubiquilin variants were cloned into pET28a vector and have the general structure of His_6_-3C-3xFlag-Ubiquilin.

For binding assays with a single substrate, the sequence of interest (Omp25 TMD, PXX region, or placeholder sequence) was cloned into the barcode 0 construct between the KpnI and BamHI sequences. The final construct had the general structure 3xHA-Sec61β soluble domain-test sequence-3F4.

The crystallization construct was cloned into a pET28b vector with N-terminal His_6_ tag followed by a 3C protease site^56^ . The fusion construct consisted of the Sti1 domain of M. bicupsidata Dsk2 (residues 165-234) followed by the receiver domain of *M. xanthus* FrzS (residues 3-124) and a variant of the Vamp2 TMD (MMIALGVACAIALAIAAVYF). The Vamp2 TMD is the same sequence used in the barcode 5 data which showed the strongest binding to Ubiquilins **(Figure S5)**. The linker between the Sti1 and FrzS domain is WGS and the linker between the FrzS domain and TMD is GNS.

### Expression and Purification of Ubiquilins

Ubiquilins were expressed in BL21 DE3 cells with the pRIL plasmid (Agilent). Cells were grown in terrific broth at 37°C until OD_600_ = 0.6-0.8. Ubiquilin protein expression was induced by addition of 0.5 mM isopropyl β-D-1-thiogalactopyranoside (IPTG, RPI corp). Upon induction, temperature was dropped to 16° C and cells were incubated for 16 h. Cells were pelleted by centrifuging for 20’ at 3900 rpm in Eppendorf 5810R centrifuge. The cell pellet from 1 L of culture was resuspended in 50 mL of UBQLN Lysis Buffer (50 mM sodium phosphate pH 7.5, 150 mM NaCl, 0.01 mM EDTA, 10% glycerol), supplemented with 0.05 mg/mL lysozyme (Simga) and 1 mM Phenylmethylsulfonyl fluoride (PMSF). Samples were stored at -80° C until purification.

Cell pellets were thawed in water and then kept on ice throughout the procedure to minimize proteolysis. After thawing, 2 μL of Universal Nuclease (Pierce) was added to each cell pellet and volume was brought to ∼120 mL by addition of lysis buffer or additional cell pellets. Cells were transferred to a metal cup surrounded by ice and lysed by sonication. The supernatant was isolated by centrifugation at 18,500 x g for 30’ at 4° C and purified by Ni-NTA affinity chromatography (Thermo Fisher) on a gravity column in the cold room. Ni-NTA resin was washed with 25 column volumes (CV) of UBQLN Lysis Buffer. Sample was eluted with Lysis Buffer supplemented with 250 mM imidazole. The eluent was concentrated to < 1 mL in a 50 kDa MWCO Amicon Ultra centrifugal filter (Millipore). Sample was then incubated overnight at 4°C with a 100:1 ratio of Ubiquilin to 3C protease to remove the His_6_ tag.

The protein was further purified by size exclusion chromatography (SEC) on a Superdex 200 Increase 10/300 GL, GE Healthcare in UBQLN FPLC Buffer (20 mM Tris pH 7.5, 100 mM NaCl). Peak fractions were pooled, concentrated to 5-15 mg/mL in a 50 kDa MWCO Amicon Ultra centrifugal filter (Millipore) and aliquots were flash-frozen in liquid nitrogen and stored at - 80°C. Protein concentrations were determined by A_280_ using a calculated extinction coefficient (Expasy).

### Expression and Purification of crystallization construct

The crystallization construct was grown and expressed as described above for Ubiquilins with the following changes. The crystallization construct was expressed in BL21 DE3 pRIL cells and grown in terrific broth until OD600 of 0.6-0.8, at which point expression was induced 0.5 mM IPTG at room temperature for 16 hours. Lysis Buffer is 50 mM Tris pH 7.5, 20 mM Imidazole, 500 mM NaCl, 10% glycerol, 1 mM DTT and Elution Buffer is 50 mM Tris pH 7.5, 500 mM imidazole, 150 mM NaCl, 1 mM DTT. After elution, the eluent was concentrated to < 1 mL in a 10 kDa MWCO Amicon Ultra centrifugal filter (Millipore). Sample was then mixed with a 100:1 ratio of protein to 3C protease and dialyzed overnight at 4° C in Dialysis Buffer (50 mM Tris pH 7.5, 150 mM NaCl, 1 mM β-mercaptoethanol). The protein was further purified by size exclusion chromatography (SEC) on a Superdex 200 Increase 10/300 GL column equilibrated in Dialysis Buffer. Peak fractions were pooled, concentrated to > 14 mg/mL in a 10 kDa MWCO Amicon Ultra centrifugal filter (Millipore) and aliquots were flash-frozen in liquid nitrogen and stored at -80°C. Protein concentrations were determined by A_280_ using a calculated extinction coefficient (Expasy).

### Protein crystallization and structure determination

Crystals of Sti1-Frzs-TMD construct were grown at 4*°* C using hanging-drop vapor diffusion method by mixing equal volume (1 µL) of protein and reservoir solution containing 2.5% MPD, 100 mM Tris pH 8.5. Large crystals were grown over a period of three days and shown to belong to the space group P12_1_1. Protein crystals were then soaked 2-3 times in cryoprotectant containing the well solution and 20% glycerol and flash frozen in liquid nitrogen.

Diffraction data were collected at Beamline 8.2.2 at Advanced Light Source at Lawrence Berkeley National Laboratory (Berkeley, CA). Data were processed and scaled using Mosflm^57^ and CCP4^58^. Crystals of Sti1-Frzs-TMD diffracted to 1.98 Å resolution and belong to space group P12_1_1 with unit cell dimensions a= 67.41 Å, b= 63.92 Å, and c= 67.43 Å. Phases were determined by molecular replacement using Phaser-MR^59^ program incorporated in the Phenix package^60^. Receiver domain from Myxococcus xanthus social motility protein FrzS (PDB: 2GKG) was used as the search model which represents 55% of Sti1-Frzs-TMD construct. The initial solution from molecular replacement generated interpretable electron density map. Model building and refinement was carried out using COOT^61^. The resulting model was further improved by positional and anisotropic B-factor refinement in Phenix, and the model quality was monitored throughout all stages of refinement process and validated using MolProbity^62^. The atomic coordinates and structure factors are deposited in the PDB: 9CKX.

### Size Exclusion Chromatography

Analytical size exclusion chromatography was performed on a BioRad NGC Quest 10 Plus with a Superdex 200 Increase 10/300 GL column (Cytiva). A total of 0.5 mL of sample at 2 mg/mL was loaded onto the column at a flow rate of 0.5 mL/min. Aldolase (158 kDa), Conalbumin (75 kDa), Ovalbumin (44 kDa), and Lysozyme (14 kDa) were used as molecular weight standards.

### Barcoded Binding Assay

In Vitro Translation (IVT) was performed using the PURExpress in vitro protein synthesis kit (NEB). Each IVT reaction contained 4 μL of PURE Solution A, 3 μL of PURE Solution B, 1 μL of EasyTag L-[^35^S] methionine stabilized aqueous solution (Perkin Elmer), 0.2 μL of Superase-in RNase inhibitor (Invitrogen), 120 ng of plasmid DNA, and a final concentration of 3 μM Ubiquilin. Nuclease free water was added to bring the total reaction volume to 10 μL. The reaction was incubated at 37° C for 2 h.

The barcoded binding assay was performed with a mixture of each of the six plasmids such that there was roughly equal expression of each construct. The ratio of each plasmid was determined empirically by running individual IVT reactions for each plasmid within a substrate series. All six reactions were run on the same SDS PAGE gel and analyzed by autoradiography and ImageJ.

After the 2 h incubation, the IVT reaction was diluted with 40 μL of Wash Buffer (20 mM Tris pH 7.5, 100mM NaCl, 1 mg/mL BSA). A 5 μL sample was taken as the “INPUT” fraction. The remaining 45 μL were added to 2 μL of settled anti-Flag magnetic beads (4 μL of 50% slurry) (Thermo Fisher) that had been equilibrated in Wash Buffer. Sample was allowed to bind to resin for 30’ at 4° C before 5 μL of sample were taken for the “FLOW THROUGH” fraction, the remaining flow through was discarded. Beads were then washed 4x with 1 mL of wash buffer at 4° C for 10’. After the final wash, 25 μL of sample were taken for the “WASH” fraction. To elute Ubiquilins, 25 μL of Wash Buffer containing 1 mg/mL 3x Flag Peptide (ApexBio) was added and the sample was incubated in the Thermomixer at 37° C for 30’. All 25 μL were taken for the “ELUTE” fraction.

The “INPUT”, “FLOW THROUGH”, “WASH”, AND “ELUTE” fractions were brought to a final volume of 25 μL by adding MilliQ Water and then 8.8 μL of 4x SDS PAGE Loading Buffer was added to the samples. Samples were then run on a 4-20% gradient Criterion gel (Bio-Rad), stained with Coomassie, and imaged.

Gel was then dried either by incubating in drying solution (40% methanol, 10% glycerol, 7.5% acetic acid) for 5’ and then loading into a gel drying apparatus (Promega) and allowed to sit at room temperature for 12-16 h with a fan on the apparatus. Alternatively, the gel was placed on filter paper and dried with a gel drier (BioRad). Dried gel was then exposed to X-ray film (RPI corp) for 24 h and then developed. Band intensity was measured using ImageJ.

Binding assays with a single substrate were carried out as described above, except the plasmid DNA for a single substrate was used. The final reaction still contained 120 ng of plasmid DNA.

### Software

The ImageJ software was downloaded here https://imagej.nih.gov/

GRAVY scores were calculated here https://www.gravy-calculator.de/

Agadir scores were calculated here http://agadir.crg.es/

**Table 1.**
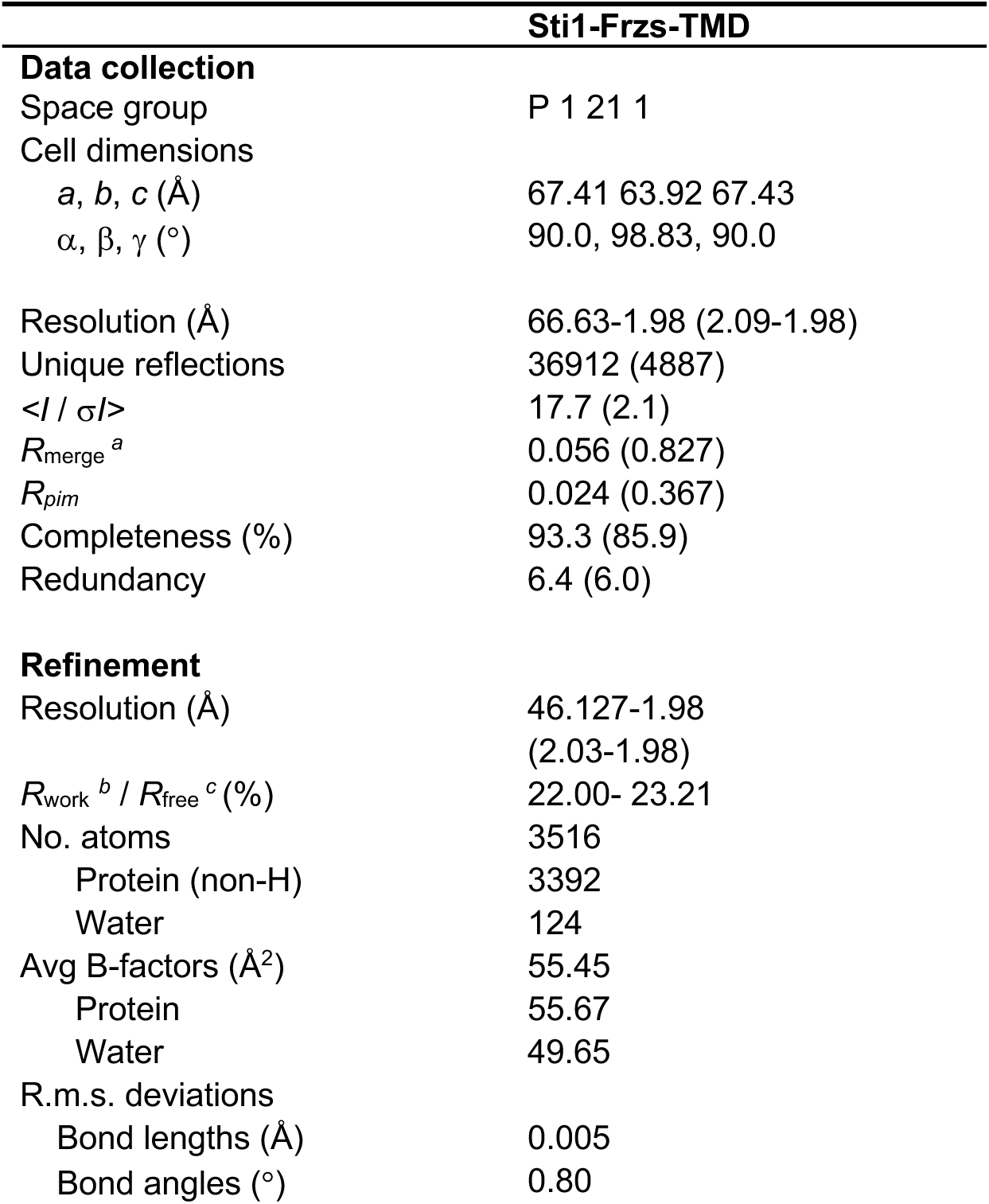
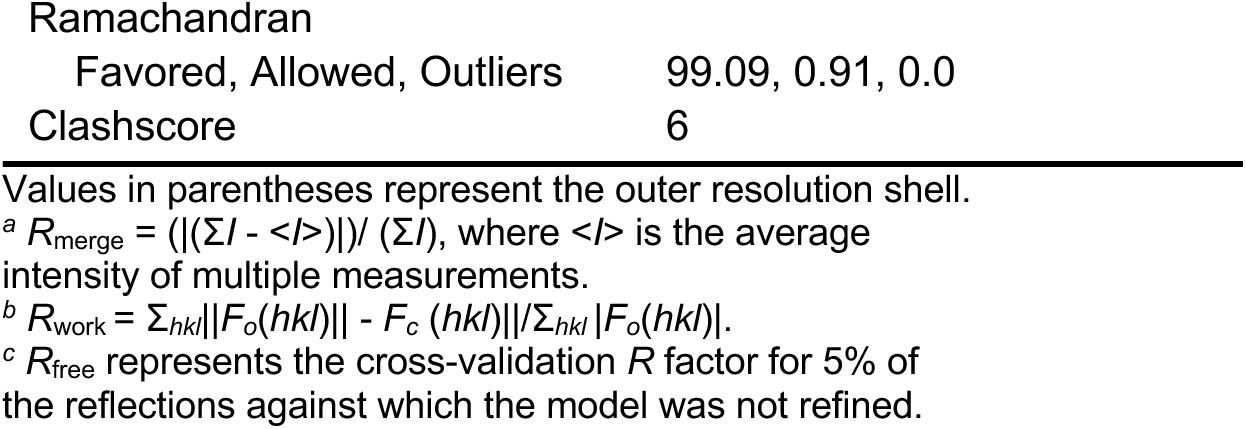
Data collection and refinement statistics.

## Acknowledgements

The authors wish to thank members of the Castañeda and Wohlever labs for helpful discussions and feedback on the project.

## Author Contributions

Conceptualization: JO, SB, SA, MLW; Data curation: JO, SB, SA, BS, MLW; Formal analysis: JO, SB, SA, MLW; Funding acquisition: MLW; Investigation: JO, SB, SA, BS; Methodology: JO, SB, SA, MLW; Project administration: MLW; Resources: JO, SB, SA, BS, MLW; Software: Not applicable; Supervision: MLW; Validation: JO, SB, SA; Visualization: JO, SB, SA, MLW; Writing – original draft: JO, MLW; Writing – review & editing: JO, SB, SA, BS, MLW;

## Supplemental Figures

**Figure S1:**
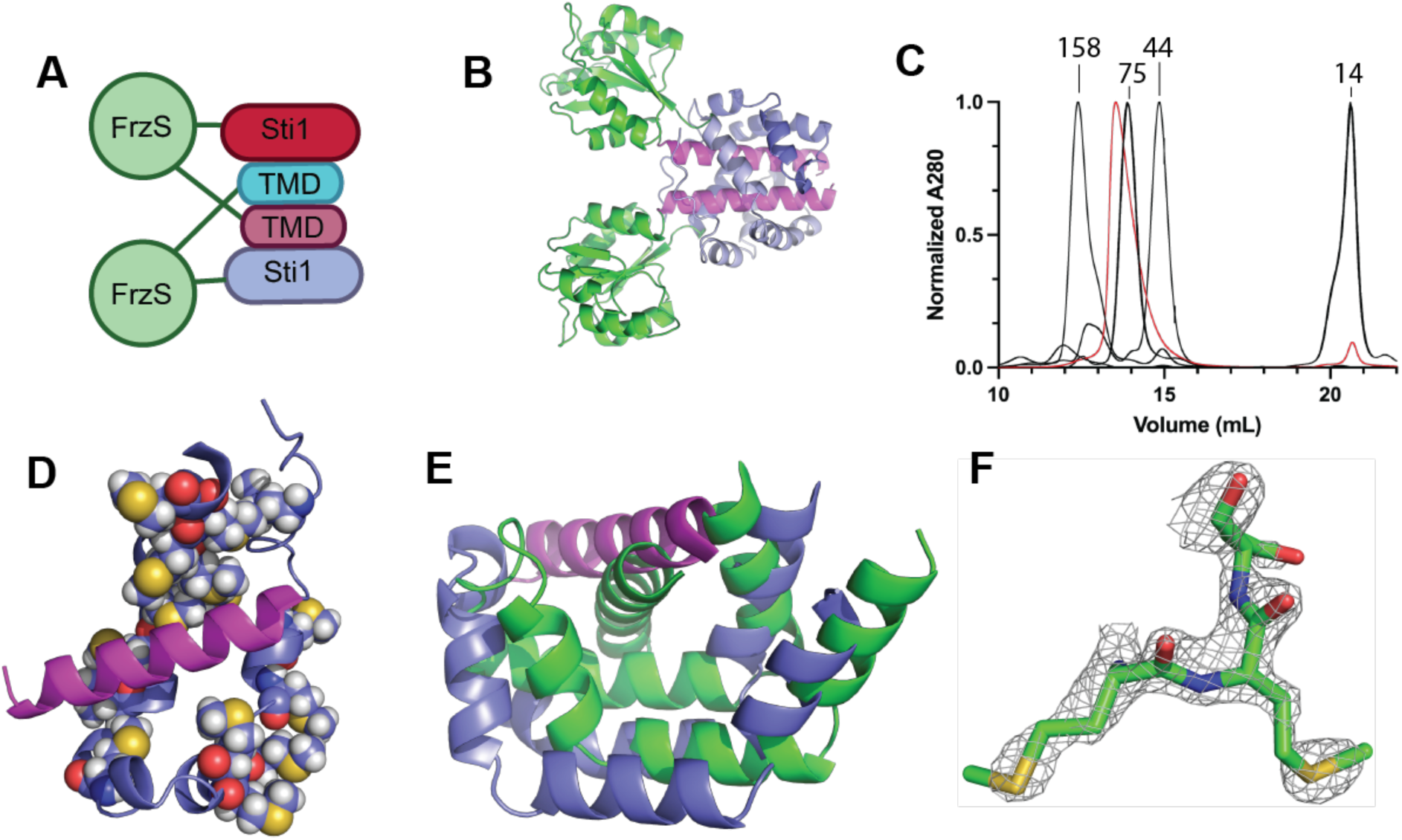
Crystal structure of the Dsk2 Sti1 domain bound to a TMD. A) Diagram showing the domain swap in the crystal structure. B) The FrzS domain does not drive dimerization and has minimal contacts with the Sti1 domain. C) Size exclusion chromatography shows that the crystallization construct (red), which has a molecular weight of 24.4 kDa, runs as an oligomer. Molecular weight of standards (Aldolase, Conalbumin, Ovalbumin, and Lysozyme) are noted on the chart. D) The hydrophobic groove is enriched for methionine residues, which are shown as spheres. E) Comparison of the crystal structure (slate and magenta) with the AlphaFold prediction (green). FrzS domains omitted for clarity. F) Fitting of the final refined model for residues 68-70 in the 0A-weighted 2*F*o-*F*c electron density map contoured at 3 sigma.

**Figure S2:**
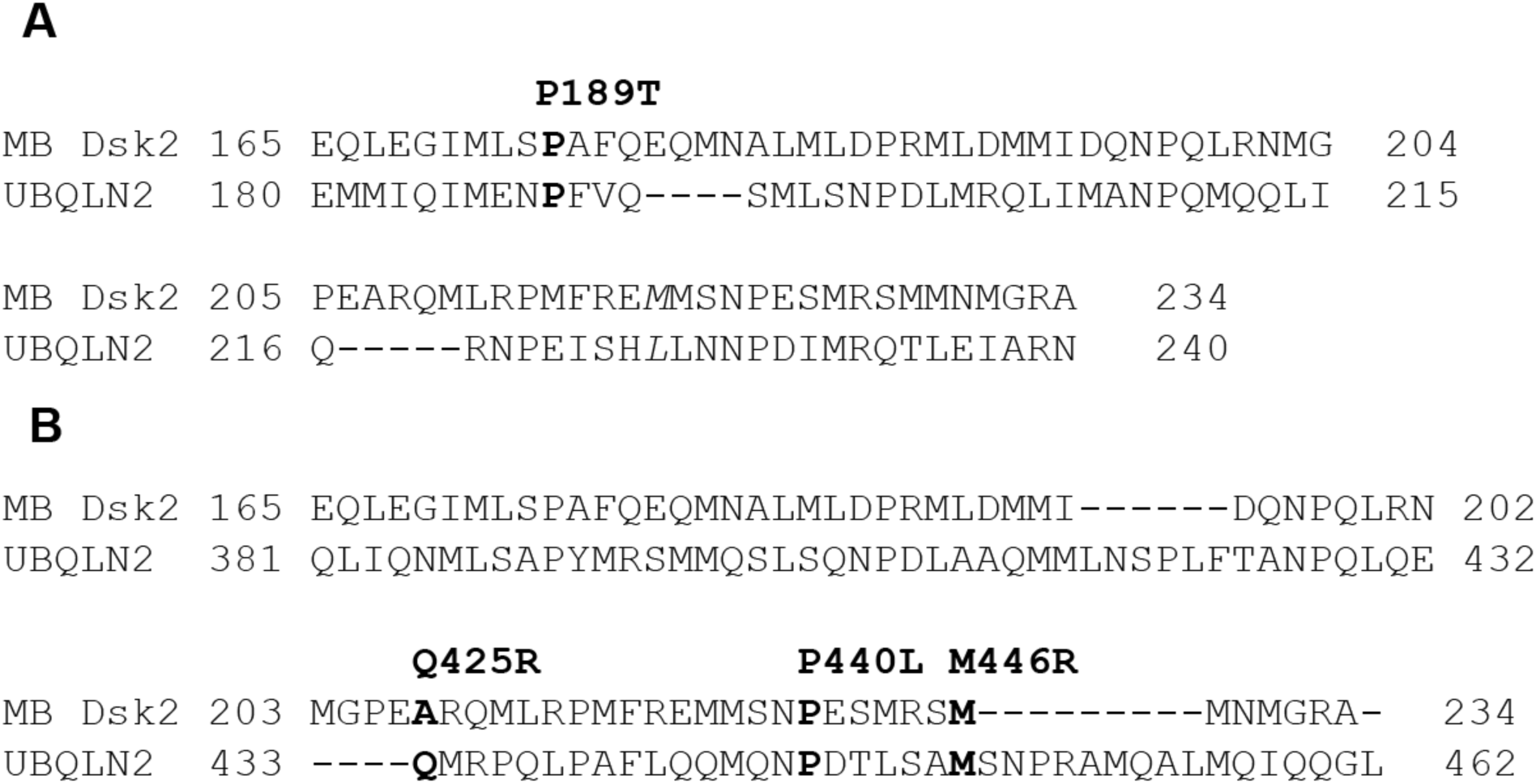
Sequence alignment of *M. bicuspidata* Dsk2 with UBQLN2. Sequence alignment between *M. bicuspidata* Dsk2 and the UBQLN2 Sti1-I **(A)** or Sti1-II **(B)** domain. Separate alignments for Sti1-I and Sti1-II were performed in Clustal Omega using the annotated domains from UBQLN1, UBQLN2, & UBQLN4. For clarity, only UBQLN2 is shown. This alignment was used to map UBQLN2 ALS mutations onto the Dsk2 structure for analysis in Figure 2.

**Figure S3:**
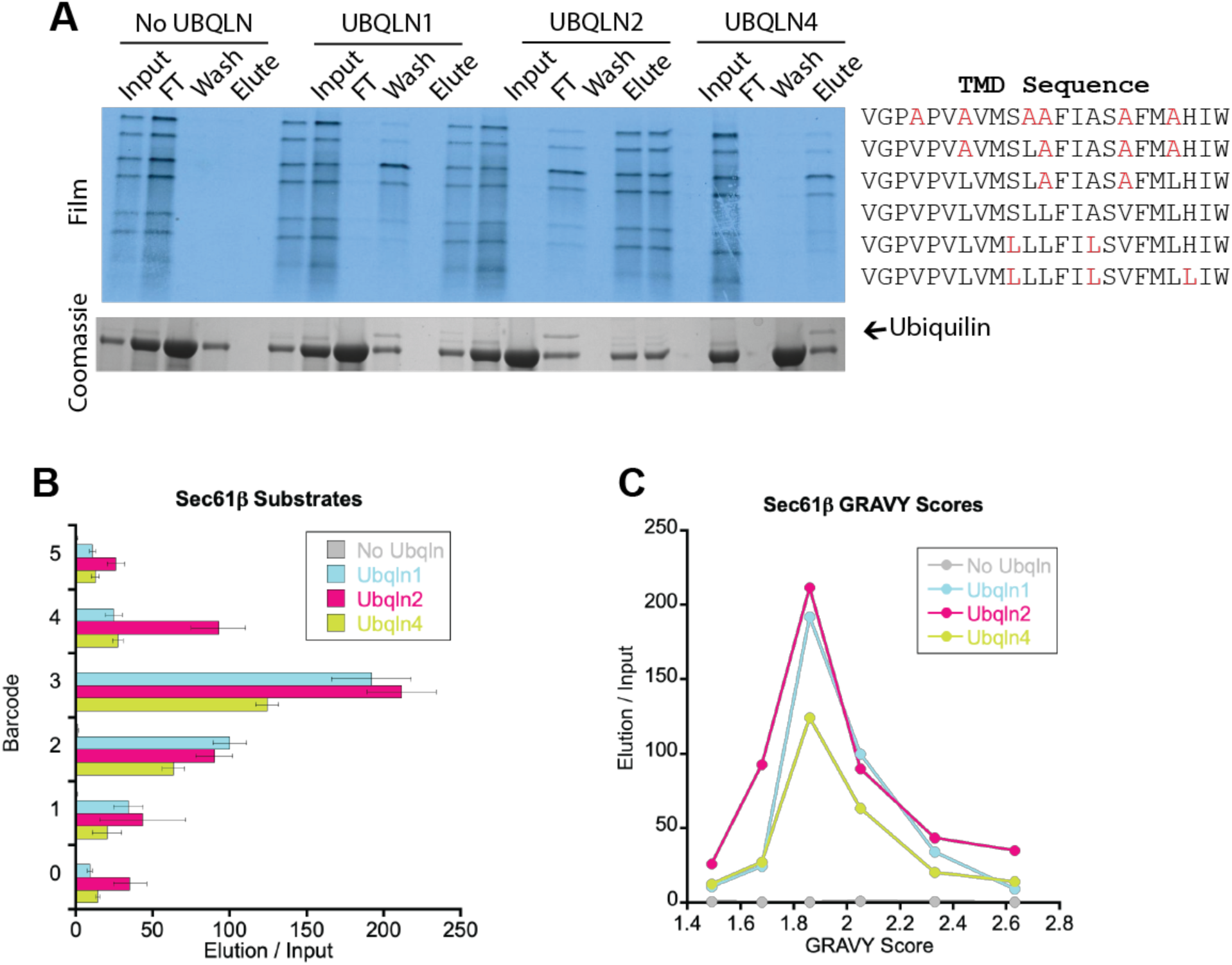
Barcoded binding data with the Sec61β substrate series. A) Barcoded binding assay with the Sec61β substrate series. The wild type Sec61β TMD has a barcode of 2. B) Quantification of the data in A, organized by barcode. A value of 100 corresponds to equal intensity of input and elution bands. C) Quantification of the data in A, organized by TMD hydrophobicity (GRAVY score). Error bars omitted for clarity.

**Figure S4:**
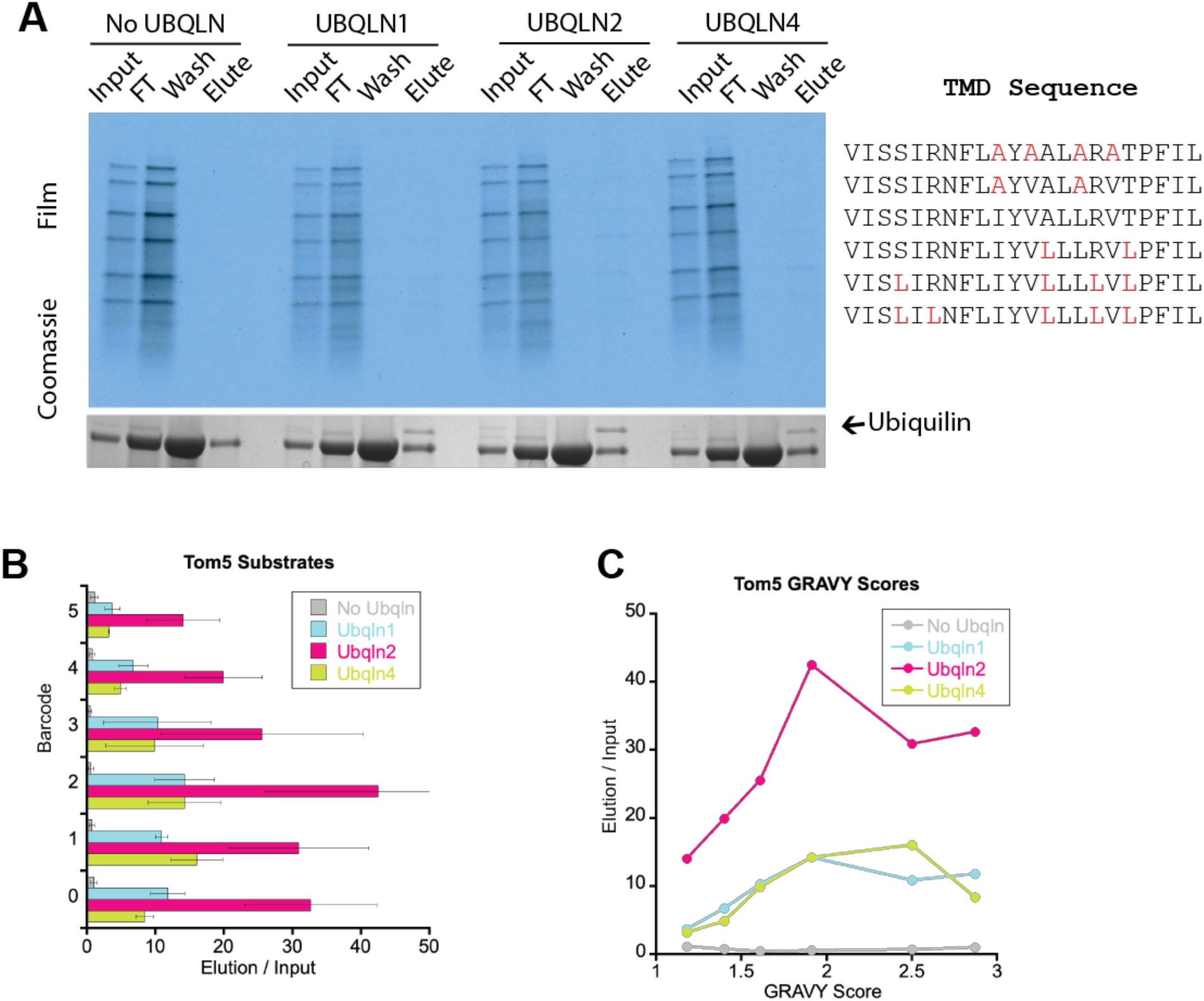
Barcoded binding data with the Tom5 substrate series. A) Barcoded binding assay with the Tom5 substrate series. The wild type Tom5 TMD has a barcode of 3. B) Quantification of the data in A, organized by barcode. A value of 100 corresponds to equal intensity of input and elution bands. C) Quantification of the data in A, organized by TMD hydrophobicity (GRAVY score). Error bars omitted for clarity.

**Figure S5:**
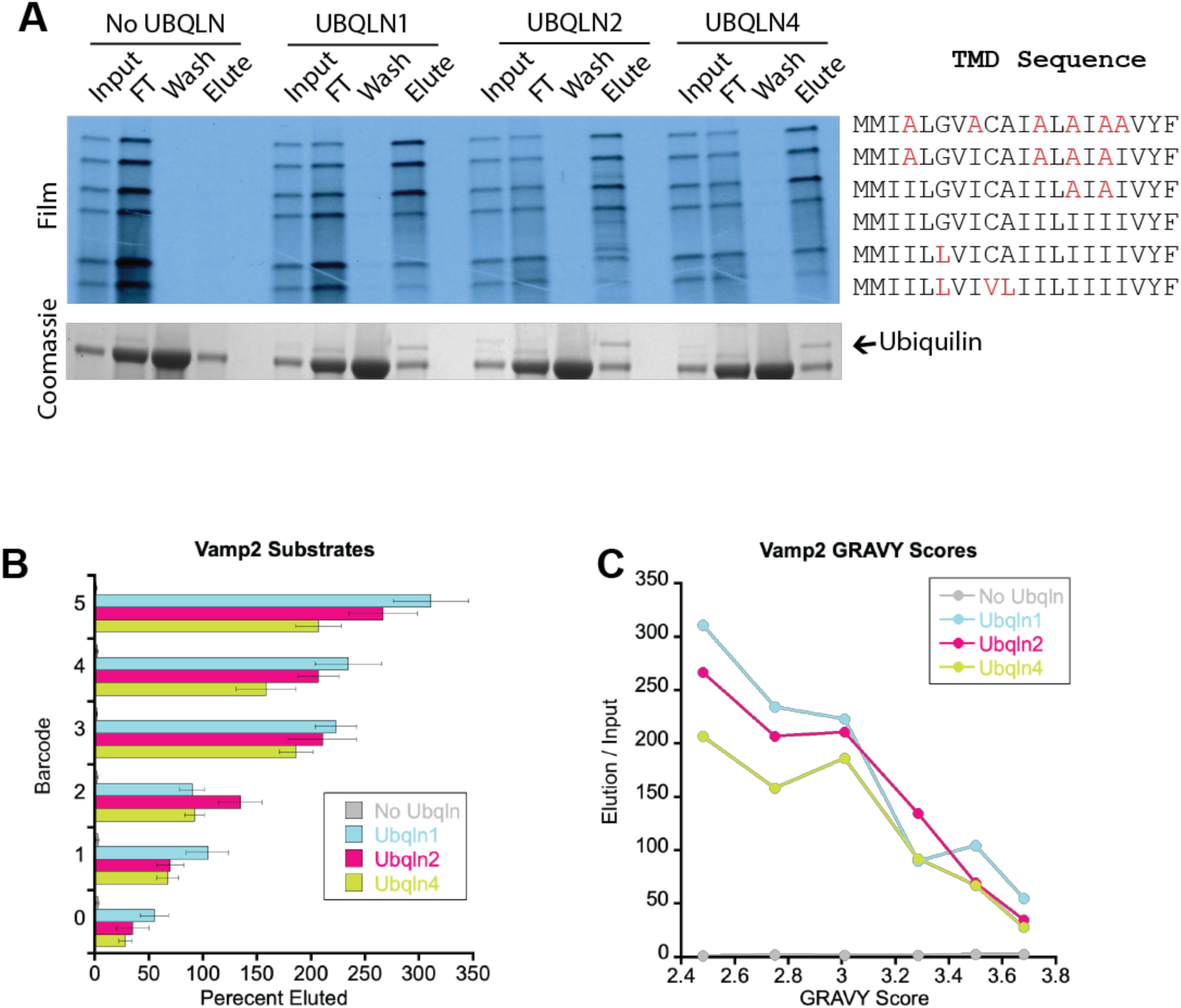
Barcoded binding data with the Vamp2 substrate series. A) Barcoded binding assay with the Vamp2 substrate series. The wild type Vamp2 TMD has a barcode of 2. B) Quantification of the data in A, organized by barcode. A value of 100 corresponds to equal intensity of input and elution bands. C) Quantification of the data in A, organized by TMD hydrophobicity (GRAVY score). Error bars omitted for clarity.

**Figure S6:**
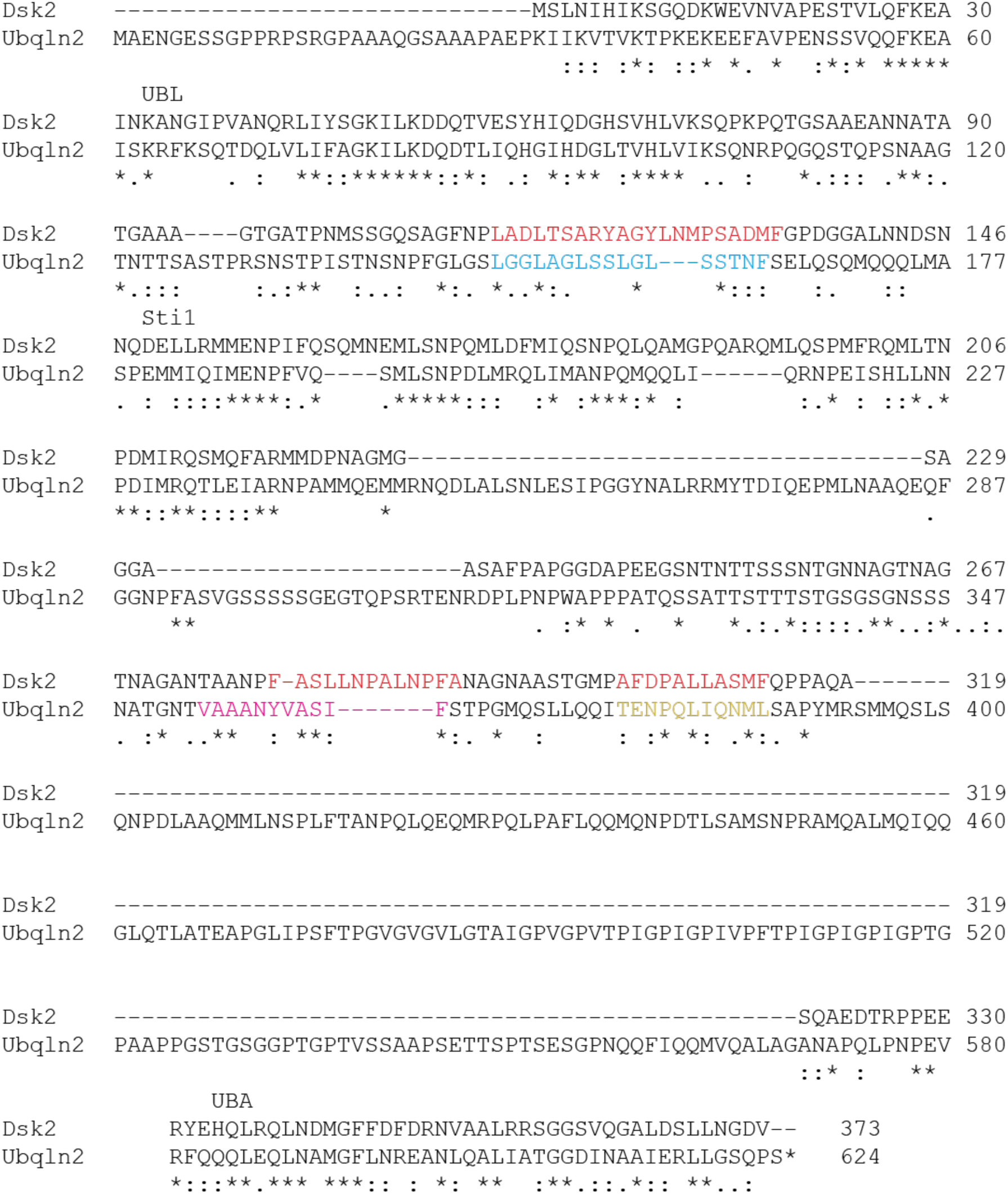
Sequence alignment show moderate conservation of placeholder sequences in Dsk2 and UBQLN2. Sequence alignment of *S. cerevisiae* Dsk2 with human UBQLN2. Dsk2 sequences highlighted in red were identified by Acharya et al. as potential Sti1 interacting motifs^53^ . UBQLN2 sequences highlighted in cyan, magenta, and yellow were chosen based on a combination of sequence alignments, AlphaFold predictions, GRAVY, and Agadir scores.

**Figure S7:**
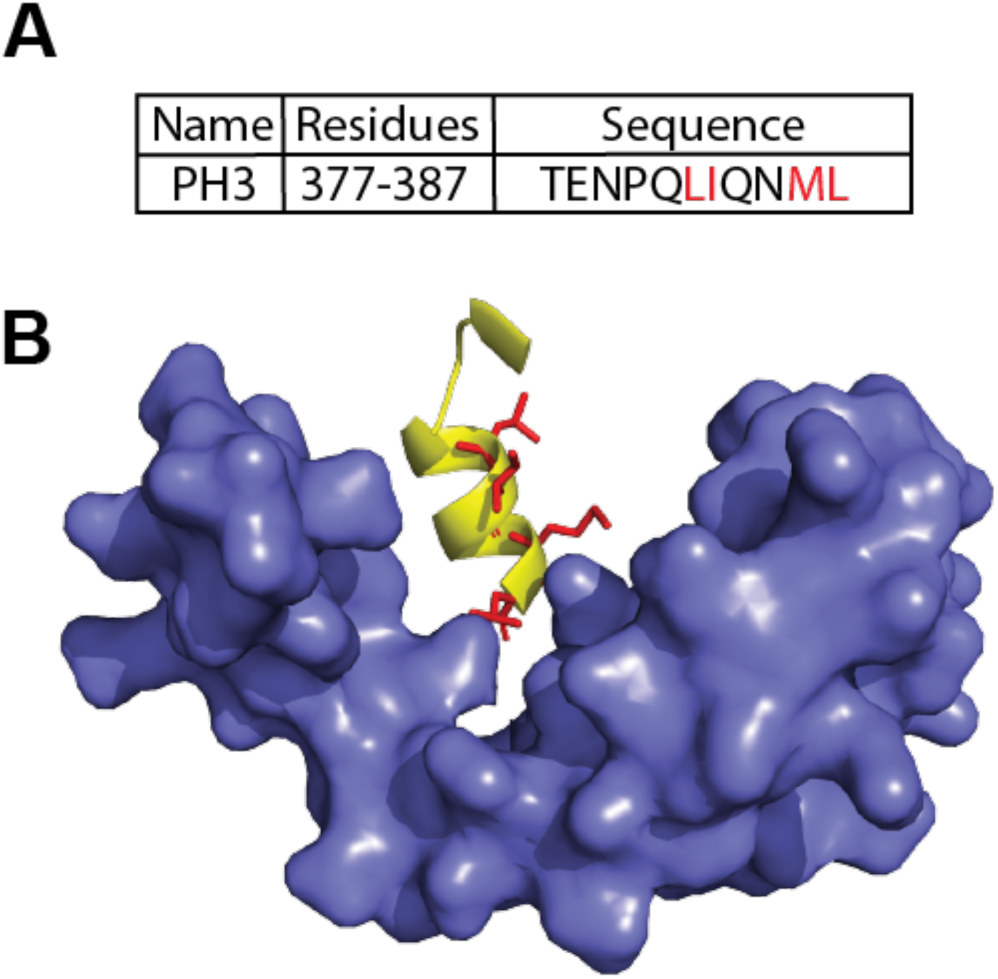
Placeholder 3 has a hydrophobic face and is positioned to interact with Sti1. A) Placeholder 3 sequence with hydrophobic residues highlighted in red. B) Alphafold 2 model of Ubiquilin 2 shows that PH3 forms an amphipathic helix with hydrophobic residues, shown in red, well positioned to interact with the hydrophobic groove of Sti1-2 surface.

